# Ultrasound transcutaneous auricular vagus nerve stimulation enhances semantic processing

**DOI:** 10.64898/2026.02.20.707135

**Authors:** Youbin Kang, Marcus Kaiser, JeYoung Jung

## Abstract

Efficient access to conceptual knowledge is central to everyday cognition, yet the effects of non-invasive neuromodulation on semantic retrieval remain insufficiently characterised. Transcutaneous auricular vagus nerve stimulation (taVNS) offers a mechanistically distinct route to modulate distributed cognitive systems via ascending vagal afferent pathways, but its effects on semantic processing has not been explored. Here, we tested whether ultrasound-based taVNS delivered to the right cymba conchae can acutely enhance semantic retrieval efficiency in healthy adults. Twenty-seven participants completed a single blind, sham-controlled study comprising two counterbalanced sessions (active vs. sham) in a within-subject design, separated by at least five days. In each session, participants performed a semantic association task and a number-judgement control task before and immediately after 30 minutes of stimulation. Active taVNS produced selective post-stimulation improvements in semantic retrieval performance relative to sham and baseline, while performance on the control task remained unchanged. Adverse effects were minimal and did not differ between active and sham conditions. These findings provide causal evidence that ultrasound-based taVNS can acutely improve the efficiency of semantic processing, expanding the cognitive profile of taVNS beyond predominantly episodic and emotional memory domains and supporting its feasibility as a scalable approach for memory enhancement.

## Introduction

Efficient access to conceptual knowledge is central to everyday activities such as communications and object use. Semantic cognition refers to the ability to use, manipulate, and generalise knowledge in order to interact effectively with the world by producing time- and context-appropriate behaviour (Binder et al., 2009; Lambon Ralph et al., 2017). This capacity supports core cognitive functions including the interpretation of word meaning, recognition of objects, and the identification of relationships between concepts, enabling rapid integration of meaning during comprehension and decision making (Binder & Desai, 2011; Lambon Ralph et al., 2017; Vatansever et al., 2017). Contemporary controlled semantic cognition (CSC) framework (Lambon Ralph et al., 2017) characterise semantic cognition not as passive knowledge storage alone, but as the dynamic interaction between representational systems and control mechanisms that shape retrieval according to current task demands and contextual constraints (Chiou et al., 2018; Jackson et al., 2021; Jefferies, 2013). Semantic cognition is supported by a complex, distributed architecture in which transmodal representational hubs within the anterior temporal lobes interact with control regions in frontal and temporoparietal cortex to regulate context- and time-appropriate retrieval (Hoffman et al., 2018; Jefferies & Wang, 2021; Lambon Ralph et al., 2017). Even in healthy individuals, variability in semantic retrieval efficiency becomes evident under conditions of time pressure, ambiguity, or weak association strength, where demands on semantic control processes increases revealing individual differences in retrieval efficiency (Hoffman, 2019; Jung & Lambon Ralph, 2023; Landin-Romero et al., 2016). Accordingly,

Transcutaneous auricular vagus nerve stimulation (taVNS) offers a mechanistically distinct approach to neuromodulation that may be well suited to influencing distributed cognitive systems (Coffman et al., 2014; Colzato & Beste, 2020; Naparstek et al., 2023). Stimulation of the auricular branch of the vagus nerve engages vagal afferent pathways projecting to the nucleus tractus solitarii, with downstream influence on brainstem and forebrain systems involving in regulating arousal, attentional allocation, and neural plasticity relevant to learning and memory (Badran et al., 2022; Frangos et al., 2015). Converging neurophysiological and neuroimaging findings suggest that through these ascending pathways, taVNS is thought to modulate large-scale network implicated in attention, emotion, cognitive control, and memory, rather than exerting focal, site-specific cortical effects (Bömmer et al., 2024; Murphy et al., 2023).

Consistent with this mechanism, a growing literature has examined the effects of taVNS on cognition, with a particular emphasis on memory-related outcomes. Most studies have focused on emotional memory, episodic memory, or associative learning (Aranberri Ruiz, 2024; Bömmer et al., 2024; Briand et al., 2020). For instance, taVNS applied during lexical decision tasks has been associated with modest improvements in subsequent high-confidence recognition responses, suggesting modulation of recollection-related memory processes (Giraudier et al., 2020). Similarly, continuous taVNS during exposure to emotionally arousing scenes has been shown to enhance long-term recognition memory, accompanied by increased parietal ERP markers of recollection at retrieval (Ventura-Bort et al., 2021). While these findings demonstrate that taVNS can influence memory performance, they primarily implicate encoding, consolidation, or emotion-related mechanisms, rather than semantic retrieval processes.

To date, most taVNS studies have relied on transcutaneous electrical stimulation of the auricular vagus nerve. However, recent developments in non-invasive neuromodulation have highlighted low-intensity focused ultrasound as an alternative means of stimulating peripheral and central neural pathways. Ultrasound-based stimulation has recently been applied to vagus nerve targets, including the auricular branch, with early work demonstrating its feasibility and physiological efficacy (Goyal et al., 2024; Ji et al., 2022). In contrast to electrical taVNS, which delivers current through the skin, ultrasound stimulation exerts its effects via mechanical interactions with neural tissue, generating localized pressure changes that can influence membrane excitability while minimising cutaneous sensations (Kaniusas et al., 2019). Initial findings suggest that ultrasound-based taVNS can influence autonomic and affective responses (Kohler et al., 2025), indicating that it may engage central vagal pathways relevant to cognitive modulation in a manner comparable to electrical approaches, while offering improved comfort and tolerability.

In the present study, we tested whether ultrasound based taVNS targeting the right cymba conchae can enhance semantic processing in healthy adults. The study employed a within subject, counterbalanced active and sham design with pre and post assessments separated by at least five days. We hypothesised that active taVNS would selectively improve semantic retrieval efficiency, compared to sham and baseline conditions, with no change on the non-semantic control task. Such findings would provide causal evidence that taVNS can modulate the efficiency of semantic retrieval dynamics, extending its cognitive profile beyond predominantly episodic and emotional memory domains.

## Methods and Materials

### Participants

27 healthy adults participated in the study (6 males; mean age = 24 ± 3.4 years; age range: 20–36 years). All participants completed a VNS Safety Questionnaire to verify eligibility. Exclusion criteria were: age under 18; presence of implanted medical devices (e.g., pacemaker, cochlear implant); history of carotid atherosclerosis or cervical vagotomy; cardiovascular conditions (including hypertension, hypotension, bradycardia, and tachycardia); metallic implants in the head or neck; current or past neurological or psychiatric disorders; tendency to faint; family history of epilepsy; use of psychoactive medication (except hormonal contraceptives); and pregnancy.

Written informed consent was obtained from all participants for both participation and publication of anonymised data. Ethical approval was granted by the University of Nottingham Ethics Committee (F1619R), and all procedures complied with the Declaration of Helsinki.

### Transcutaneous auricular vagus nerve stimulation (taVNS)

taVNS was administered using ultrasound stimulation (ZenBud, NeurGear Inc., Rochester, NY, USA), targeting the right cymba conchae. The stimulation protocol employed a centre frequency of 5.3 MHz, a pulse repetition frequency of 41 Hz, a 50% duty cycle, a peak mechanical output of 0.081 W/cm^2^, and an average pressure amplitude of 1.03 MPa.

### Experimental design and procedures

Participants completed two sessions, an active stimulation session and a sham session separated by a minimum of five days. Session order was counterbalanced across the participants. At the beginning of each session, participants performed a semantic memory task and a number judgement task as a control task. After the pre-stimulation session, participants received 30 minutes of taVNS. Immediately following stimulation (post stimulation session), both tasks were repeated (Fig. 1A). At the end of each session, participants completed a side-effects questionnaire assessing any mild adverse sensations (e.g., headache, neck tension, nausea, tingling, or ear pain).

**Figure 1.**
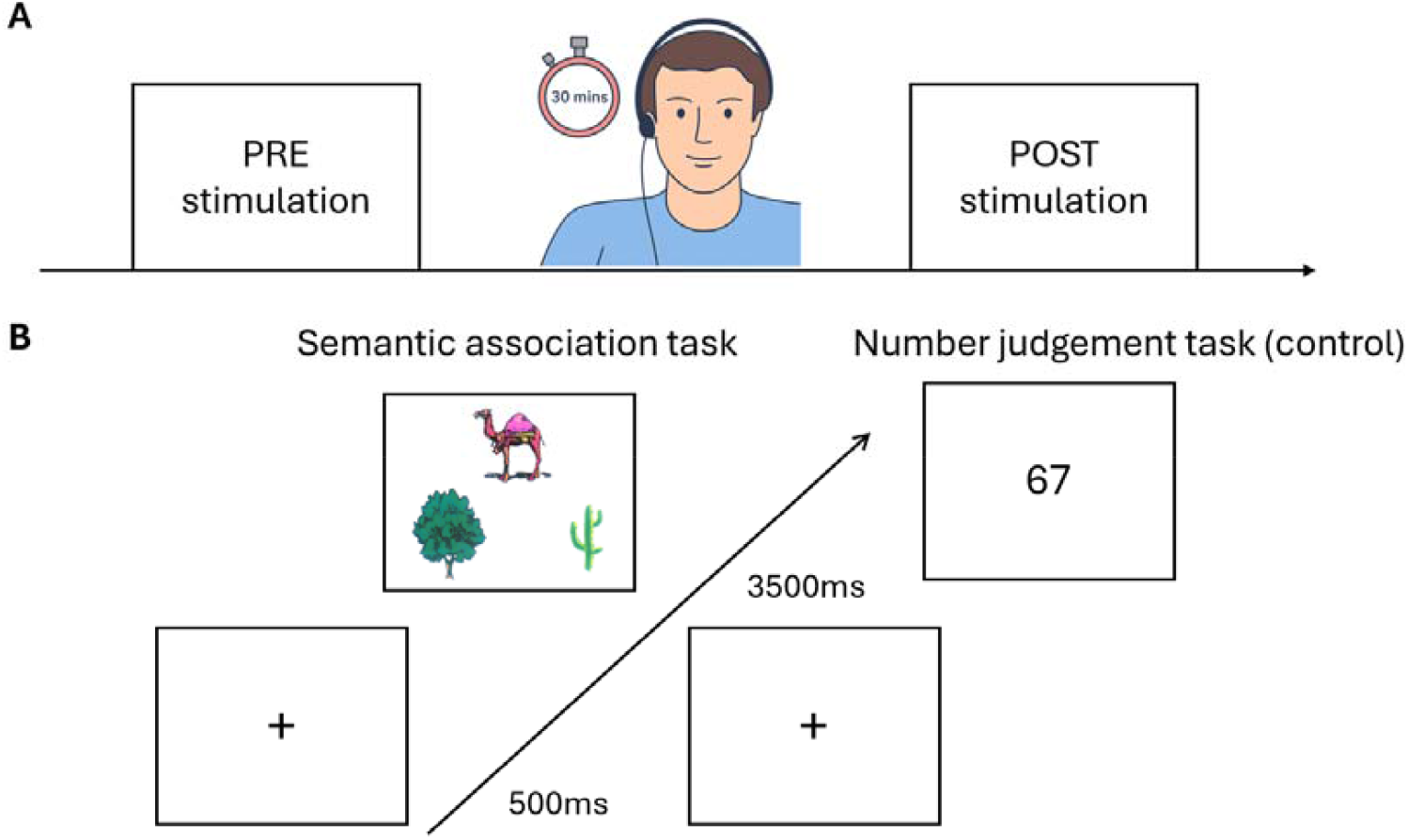
Experimental design and task strcuture. (A) Schematic of the within-subjects design. Each participant completed two sessions (active taVNS and sham), separated by at least five days, with session order counterbalanced. In each session, participants first performed the semantic association task and the number judgement task (pre-stimulation). They then received 30 minutes of taVNS or sham stimulation. Immediately after stimulation, participants completed both tasks again (post-stimulation). (B) Example trial structure for the semantic association task (left). Each trial began with a 500ms fixation cross followed by a 3500ms display of a probe image at the top and two choice images at the bottom. Participants selected the image most semantically related to the probe (e.g., camel – cactus). Example trial structure for the number judgement task (right). Following a fixation period, a randomly generated two-digit number appeared at the centre of the screen. Participants indicated whether the number was odd or even using designated response keys.

The semantic association task required participants to select which of two images presented at the bottom of the screen was semantically related to a probe image shown at the top (Fig. 1B). Stimuli were taken from the Pyramids and Palm Tree Test (Howard & Patterson, 1992) and the Camel and Cactus Test (Bozeat et al., 2000). Each trial began with a 500ms fixation cross, followed by a 3500ms presentation of the stimulus array. In the number judgement task, participants determined whether acentrally displayed two-digit number was odd or even (Fig. 1B). After a fixation period, a randomly generated number appeared, and participants responded using the same two-button setup (e.g., left key for “even,” right key for “odd”). Both tasks comprised 60 trials per session, and distinct stimulus sets were used across sessions to avoid repetition or carryover effects. All experimental tasks were created and presented using PsychoPy (version 2024.2.4).

### Statistical analysis

A within-subjects design was used, with stimulation (active vs. sham) and session (pre vs. post) as repeated-measures factors. A 2 × 2 repeated-measures ANOVA assessed the main and interaction effects of stimulation and session for each task. Planned paired-samples *t*-tests further examined pre–post changes within each stimulation condition and compared post-stimulation performance between active and sham sessions. A chi-square test was used to compare the frequency of reported side effects between active and sham stimulation. All analyses were conducted in SPSS (version 29).

## Results

### Semantic association task results

For accuracy, the 2 × 2 repeated-measures ANOVA showed no significant main effects of stimulation or session, and no interaction (stimulation: F(1,26) = 2.16, p =.153; session: F(1,26) = 0.92, p =.348; interaction: F(1,26) = 2.80, p =.106). However, planned post hoc tests revealed increased accuracy following active compared to sham stimulation in the post-stimulation session (t(26) = 2.29, p =.015, one-tailed) (Fig. 2A). For reaction time (RT), there was a significant main effect of session (F(1,26) = 4.78, p =.038), with no significant stimulation or interaction effects (ps >.34). Post hoc tests showed faster RTs after active stimulation relative to pre-stimulation (t(26) = 2.56, p =.017, two-tailed) (Fig. 2B). No other comparisons were significant (all ps >.069).

**Figure 2.**
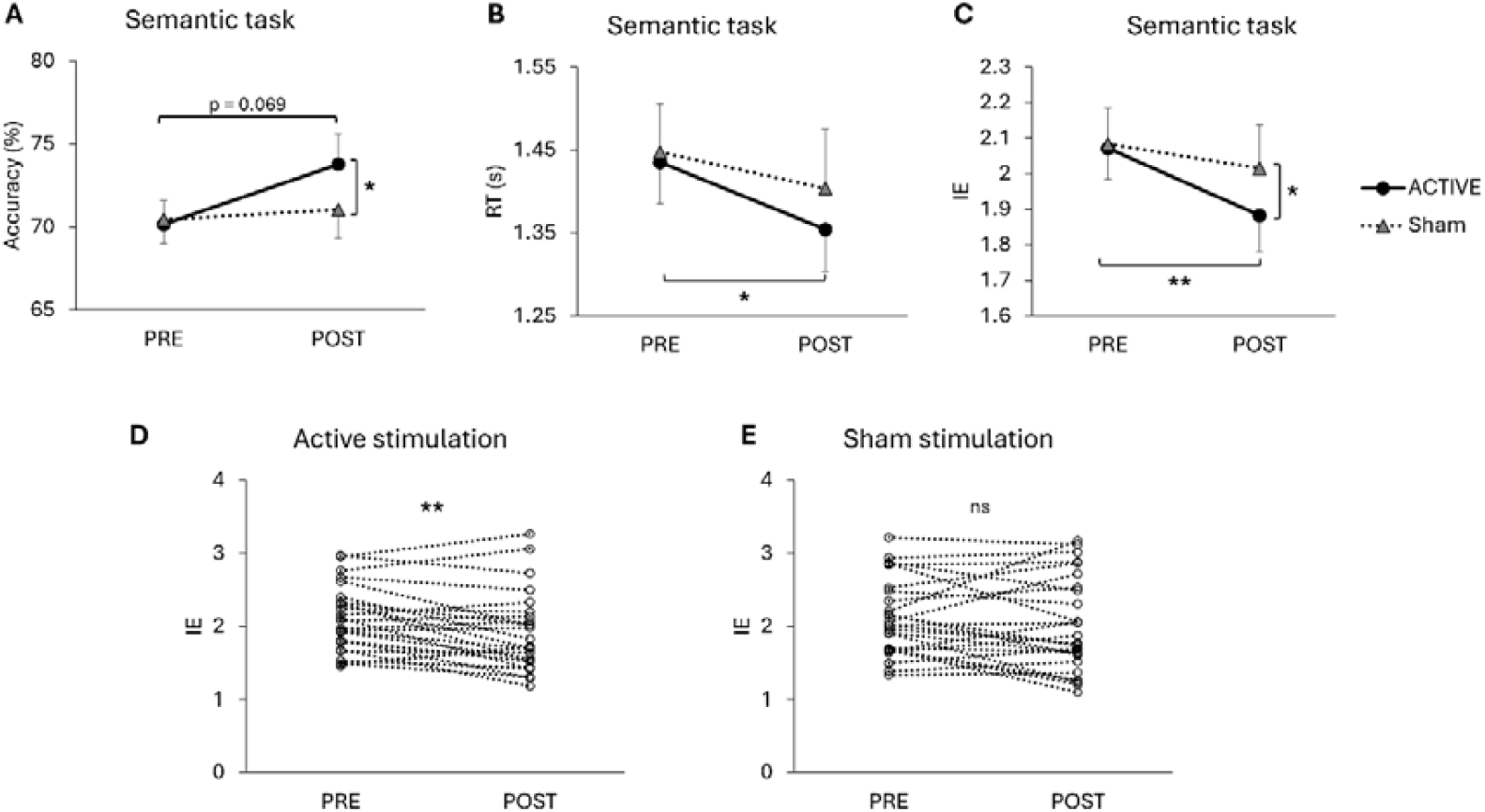
Effects of taVNS on semantic association performance. (A) Mean accuracy (± SEM) for active and sham stimulation across pre- and post-stimulation sessions. (B) Mean reaction time (RT; ± SEM) across conditions. (C) Inverse efficiency (IE = RT/accuracy × 100; higher values indicate poorer performance) for each stimulation condition and session. (D–E) Individual participant data for active stimulation (D) and sham stimulation (E), illustrating within-subject changes across sessions. Black circles and solid lines represent active stimulation, whereas grey triangles and dotted lines indicate sham stimulation. Error bars represent ± SEM. * p < 0.05, ** p < 0.01. ns = not significant.

To capture combined speed–accuracy performance, inverse efficiency (IE = RT/accuracy × 100) was computed. The ANOVA revealed a significant main effect of session (F(1,26) = 4.33, p =.047), with no significant stimulation or interaction effects (ps >.093). Planned comparisons showed that active stimulation improved performance, reflected in lower IE scores compared to both pre-stimulation (t(26) = 3.07, p =.005, two-tailed) and post-sham stimulation (t(26) = –1.72, p =.049, one-tailed) (Fig. 2C). Individual participant data are displayed in Fig. 2D–E.

### Control task results

The 2 × 2 repeated-measures ANOVA showed no significant effects for accuracy (stimulation: F(1,26) = 0.26, p =.613; session: F(1,26) = 0.11, p =.747; interaction: F(1,26) = 2.53, p =.124), RT (stimulation: F(1,26) = 0.001, p =.975; session: F(1,26) = 2.99, p =.096; interaction: F(1,26) = 1.50, p =.231), or IE (stimulation: F(1,26) = 0.02, p =.884; session: F(1,26) = 2.41, p =.133; interaction: F(1,26) = 0.15, p =.707). Planned post hoc tests showed no significant differences for any comparison (all ps >.064). Results are summarised in Fig. 3.

**Figure 3.**
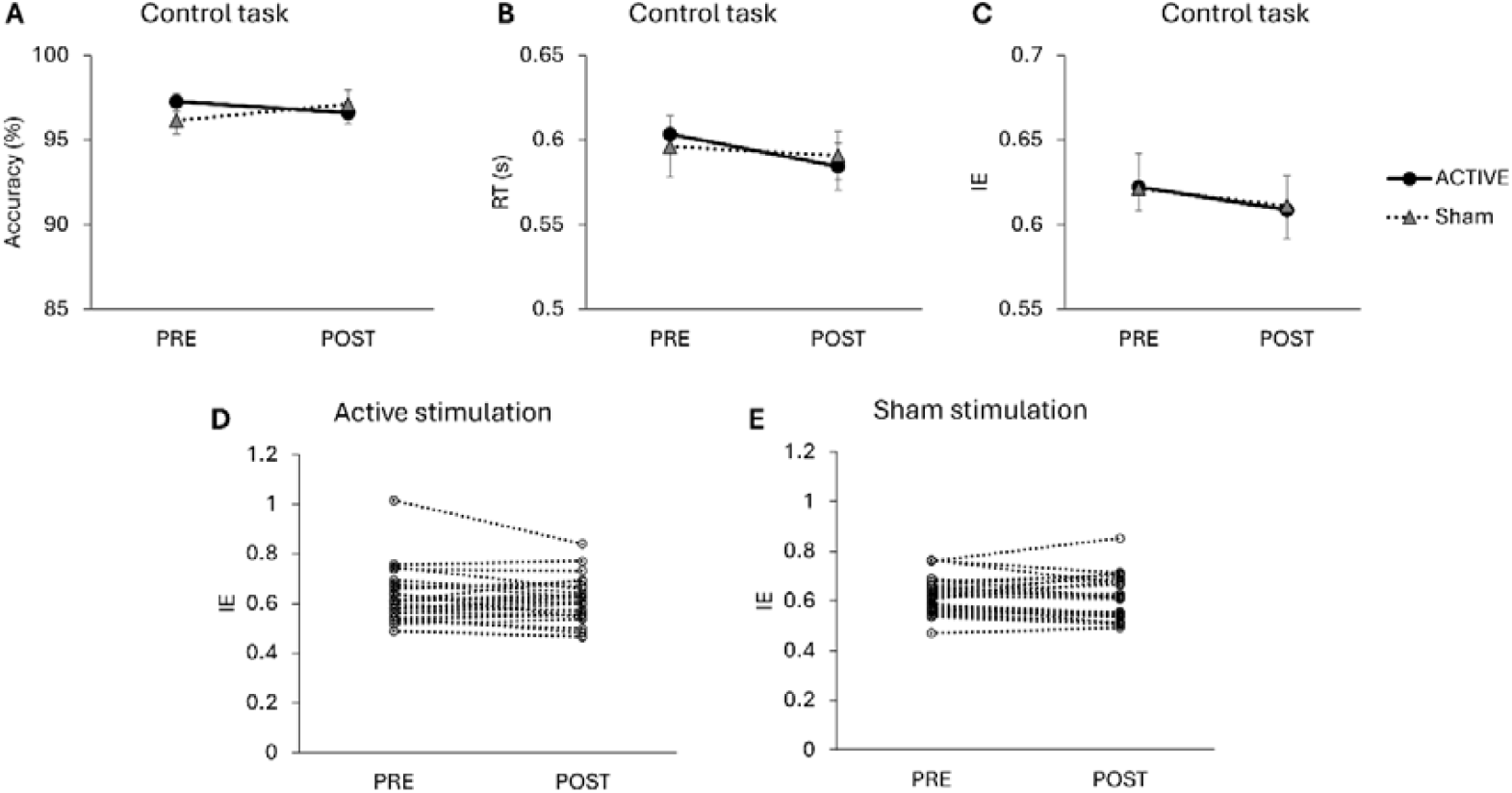
Control task performance across stimulation conditions. Mean accuracy (A), reaction time (RT; B), and inverse efficiency (IE; C) for the number judgement control task across pre- and post-stimulation sessions. (D–E) Individual participant data for active stimulation (D) and sham stimulation (E), illustrating within-subject changes across sessions. No significant effects of stimulation, session, or their interaction were observed for any measure. Error bars represent ± SEM. Black circles and solid lines represent active stimulation, whereas grey triangles and dotted lines indicate sham stimulation.

### Aversive questionnaire results

No participants reported experiencing discomfort during the study and were able to differentiate between active and sham stimulation. Questionnaire analysis showed no significant difference between active and sham stimulation (x^2^ < 1.02, ps >.313) (Table 1).

**Table 1.**
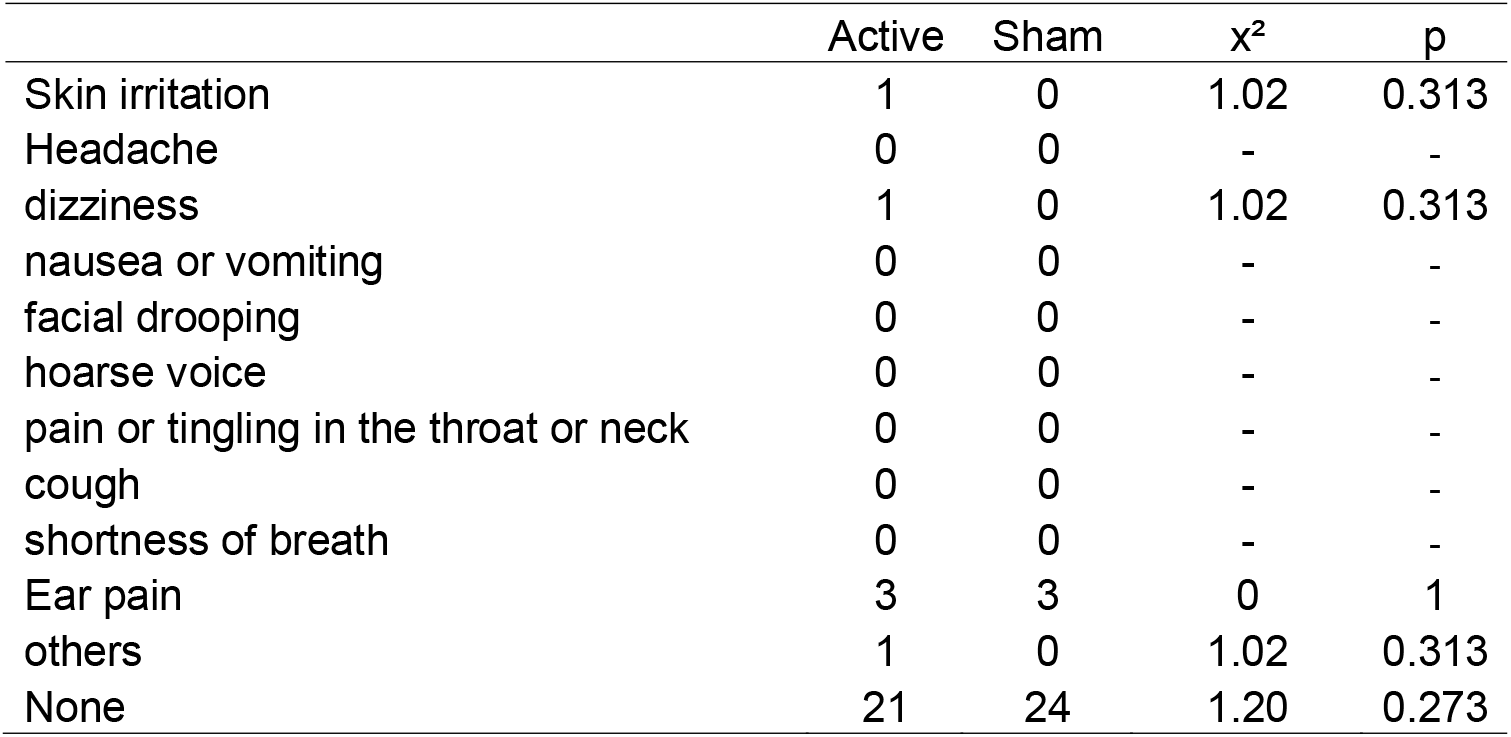
Summary of reported side effects on the aversive questionnaire. Values indicate the number of occurrences.

## Discussion

The present study examined whether ultrasound-based transcutaneous auricular vagus nerve stimulation (taVNS) can acutely modulate semantic processing in healthy adults. We found that active taVNS improved semantic association performance. Performance on a control task remained unchanged, and adverse effects were minimal and comparable between active and sham sessions. Together, these findings provide initial evidence that ultrasound-based taVNS can selectively enhance the efficiency of semantic processing.

We observed task-specific effects of taVNS, such that active stimulation selectively enhanced semantic task performance relative to sham and baseline conditions, while performance on the control task remained unchanged. The absence of change on the nonsemantic control task argues against explanations based on indiscriminate response speeding, global arousal, or motivational enhancement (Colzato & Beste, 2020). Instead, the selective improvement in semantic association performance is most consistent with improved efficiency in processes that are disproportionately taxed by semantic decision, namely rapid access to conceptual relations and controlled selection among competing alternatives under time pressure (Davey et al., 2015; Noonan et al., 2013). This interpretation aligns closely with contemporary accounts of semantic cognition, CSC, which conceptualise behaviour as emerging from interactions between distributed representational systems and semantic control mechanisms that regulate retrieval in a context-sensitive manner, particularly when time pressure amplifies competition among alternatives (Hoffman, 2019; Jefferies & Wang, 2021; Lambon Ralph et al., 2017).

At the neural systems level, the present findings are consistent with growing evidence from fMRI studies showing that taVNS modulates activity across distributed brain networks (Bömmer et al., 2024; Colzato & Beste, 2020; Frangos et al., 2015). Importantly, many of the regions influenced by taVNS including prefrontal cortex, temporal regions, angular gyrus, cingulate cortex, and limbic structures overlap with the broader semantic network supporting representational and control processes (Binder & Desai, 2011; Chiou et al., 2023; Lambon Ralph et al., 2017; Murphy et al., 2023). Rather than exerting focal effects on isolated cortical targets, taVNS appears to influence large-scale network dynamics via ascending vagal afferent pathways. Such widespread modulation is well aligned with contemporary models of semantic cognition, which emphasise interactions between anterior temporal representational hubs and fronto-temporo-parietal control systems (Hoffman et al., 2018; Jefferies & Wang, 2021; Lambon Ralph et al., 2017). From this perspective, taVNS may bias network states in a manner that facilitates more efficient coordination between these systems, thereby improving the speed and reliability with which task-relevant conceptual information is accessed and selected (Colzato & Beste, 2020; Naparstek et al., 2023; Rodenkirch et al., 2022). The present behavioural results provide functional support for this network-level account by demonstrating that stimulation selectively benefits semantic decisions that depend on distributed system interactions.

Electrophysiological evidence further supports a mechanistic link between taVNS and semantic retrieval efficiency. Across encoding and retrieval contexts, successful access to semantic information has been robustly associated with modulation of alpha-band oscillatory activity (8–13 Hz) in neocortical regions (Branzi et al., 2023; Gastaldon et al., 2020; Hanslmayr et al., 2016; Hanslmayr et al., 2012; Roos & Piai, 2020). Alpha desynchronisation is widely interpreted as reflecting increased information-processing capacity and more flexible access to stored representations, particularly under conditions of high selection demand (Hanslmayr et al., 2016; Hanslmayr et al., 2012; Jensen & Mazaheri, 2010). Emerging EEG studies indicate that taVNS can modulate alpha-band dynamics, raising the possibility that stimulation-induced changes in oscillatory state support more efficient semantic retrieval (Lloyd et al., 2023; Sharon et al., 2021; Skora et al., 2024). taVNS may enhance semantic performance by biasing networks toward states that favour rapid information integration and competition resolution.

An important contribution of the present work is the use of ultrasound-based taVNS as an alternative to conventional electrical stimulation. Unlike electrical taVNS, which delivers current through the skin and can be associated with cutaneous discomfort, ultrasound stimulation operates via mechanical interactions with neural tissue, generating local pressure changes that influence membrane excitability while minimising sensory side effects (Goyal et al., 2024; Ji et al., 2022; Kohler et al., 2025). Consistent with this, ultrasound taVNS was well tolerated in the present study, with adverse effects comparable between active and sham conditions. Beyond tolerability, ultrasound-based taVNS offers practical and translational advantages. Its focality and parameter flexibility may enable more precise engagement of vagal pathways, while its compatibility with earbud-style delivery supports scalability and repeated use. The present findings therefore demonstrate not only the cognitive relevance of taVNS for semantic processing, but also the feasibility of ultrasound-based stimulation as a practical neuromodulatory tool for targeting distributed cognitive systems.

Several limitations of the current study warrant consideration. First, the sample size and single-session protocol limit inference about individual variability, durability, and dose-response relationships. Future work should explicitly characterize the time course of the behavioral benefit by testing whether semantic-efficiency gains persist beyond the immediate post-stimulation window (e.g., at delayed follow-ups such as approximately 1 hour and 24 hours post-stimulation), and by testing whether effects accumulate or stabilize with repeated sessions. Second, although the control task reduces interpretive ambiguity, it cannot fully exclude broader state changes that disproportionately benefit semantically rich decisions. Incorporating physiological indices (e.g., pupil-linked arousal, HRV) and/or neural readouts (EEG/fMRI markers of semantic control and competition) would substantially strengthen mechanistic claims. Finally, systematic exploration of stimulation parameters including intensity, timing, and duration will be essential for establishing reproducible protocols and identifying boundary conditions for semantic enhancement.

In summary, ultrasound-based taVNS delivered to the right cymba conchae produced a selective improvement in semantic association efficiency, with no corresponding change in a control task. These findings broaden the cognitive profile of taVNS beyond predominantly episodic/emotional memory outcomes and provide initial causal evidence that vagal-afferent stimulation can enhance the efficiency of semantic cognition. The results motivate mechanistically informed follow-up studies that combine scalable stimulation delivery with physiological and neural readouts to determine when, for whom, and through which control-related pathways semantic benefits emerge.

## Author Contributions

Conceptualization: JJ; Methodology: YK, JJ, MK; Investigation: YK, JJ; Writing: YK, JJ, MK; Writing – review&editing: YK, JJ, MK.

